# Metalloprotease Gp63 targeting novel glycoside exhibits potential antileishmanial activity

**DOI:** 10.1101/2020.09.11.292920

**Authors:** Amrita Chakrabarti, Chintam Narayana, Nishant Joshi, Swati Garg, Lalit Garg, Ram Sagar, Soumya Pati, Shailja Singh

**Affiliations:** Department of Life Sciences, School of Natural Sciences, Shiv Nadar University. Uttar Pradesh 201314, India; Department of Chemistry, School of Natural Sciences, Shiv Nadar University, Uttar Pradesh 201314, India; Gene Regulation Laboratory, National Institute of Immunology, New Delhi, India; Department of Chemistry, Institute of Science, Banaras Hindu University, Varanasi 221005, Uttar Pradesh, India; Special Centre for Molecular Medicine, Jawaharlal Nehru University, New Delhi, India

**Keywords:** Leishmania, Glycoside 2, LdGp63, CETSA

## Abstract

Visceral Leishmaniasis (VL) and its aggressive cutaneous exacerbation known as Post Kala-azar Dermal Leishmaniasis (PKDL) cause a huge disease burden in tropics and sub-tropic endemic zones worldwide. Contemporary treatment modalities have been associated with various complications. Encouraged from the recent marked antimalarial effects from plant derived glycosides; here we have chemically synthesized a library of diverse Glycoside derivatives (Gly 1-12) and evaluated their inhibitory efficacy against *Ag83* strain of *Leishmania donovani. In vitro* activity of Glycoside-2 **(Gly 2)** on promastigote form of *Ag83* strain, unravelled its prominent anti-leishmanial property with **IC50** value of 1.13μM. *In-silico* studies also unveiled the efficacy of **Gly 2** to bind to the membrane surface of parasite. The toxic effect of **Gly 2** causes necrosis like death in promastigote by abrogating its proliferation leading to imbalanced redox homeostasis by disruption of mitochondrial membrane potential. Additionally, **Gly 2** treatment demonstrated increased susceptibility of parasites towards complement mediated lysis and displayed strong lethal effect on amastigote-macrophage infection model mimicking pathophysiological condition of body. This lead molecule was quite effective against the clinical on promastigotes form of PKDL strain BS12 with IC50 value of 1.97 μM making it the most suitable drug so far which can target both VL and PKDL simultaneously. Based on the above experimental validations we narrowed our thoughts regarding the potent role of **Gly 2** targeting surface protein of *L. donovani* such as Gp63, a zinc metalloprotease. Further analysis of structure activity relationship (SAR) of these glycoside derivatives, demonstrated exceptional binding affinity of **Gly 2** towards Gp63, a zinc metalloprotease of *L. donovani*; with strong H-bond interactions of **Gly 2** with catalytic domain in the α-helix B region of Gp63. The strong confined interactions between **Gly 2** and the target protein Gp63 in a physiologically relevant cellular environment was further assessed by Cellular Thermal Shift Assay **(CETSA)** which corroborated with our previous results. Taken together, this study reports the serendipitous discovery of glycoside derivative **Gly 2** with enhanced leishmanicidal activity and proves to be novel chemotherapeutic prototype against VL and PKDL.

**Highlights:**

- A novel glycoside derivative (Gly 2) targets Gp63 functioning in *L. donovani* promastigotes, resulting in its abrogated proliferation and severely detabilized redox homeostasis, leading to parasitic death.
- Structure activity relationship (SAR) analysis revealed exceptional ligandability of Gly 2 towards Gp63 catalytic domain both *in silico* and in Cellular Thermal Shift Assay (CETSA) based *in vitro* analysis.
- Gly 2 treatment exhibited increased parasite susceptibility towards complement mediated lysis and reduced macrophage infection *in vitro* mimicking the pathophysiological conditions.
- Gly 2 showed profound antileishmanial activity against the clinical isolates of Post Kala-azar Dermal Leishmaniasis (PKDL).

## 1. Introduction

Leishmaniasis is a globally neglected vector-borne disease, transmitted by the bite of female sand fly infected with the protozoan parasite of genus *Leishmania*. It has a diverse range of clinical manifestations ranging from a mild cutaneous variant to the life-threatening visceral form [1]. Those patients who are apparently cured of Visceral Leishmaniasis (VL), are prone to Post kala-azar dermal leishmaniasis (PKDL), an aggressive clinical manifestation of VL [2]. Kala-azar is caused by *L. donovani* which is endemic to several parts of the tropics, subtropics, and southern Europe; is a severe threaten to 350 million people, with a prevalence of 12 million people worldwide [3]. In addition to it, an estimated 0·7–1 million newly reported cases of leishmaniasis erupt out every year [4]. Thus, targeting the fatal disease; WHO has already declared the development of new treatments with utmost priority.

Over several years, the available diagnosis and treatment modalities for VL relied on pentavalent antimonials using compounds like sodium stibogluconate and meglumine antimoniate. But they are found to be unsatisfactory in combating the alarming situation of arising new cases leading to sudden increase in disease burden. The underlying barricades towards their failure were high toxicity, delayed drug release and response; inimical side effects and development of resistance towards the available drug regimen [5] [6]. One of the contemporary approaches includes the use of amphotericin B accompanied by emanation of miltefosine as the first oral drug therapy for VL [7]. In the conquest for identifying better leishmanicidal compounds, plant-derived and/or mimicking products have been gaining ground. This includes Aloe vera, luteolin, quassin, berberine chloride and artemisinin among others [8] which share a common mode of action; inducing oxidative stress leading to parasite death. But they too depicted certain side-effects and low efficacy. Thus, briefly current armamentarium of anti-leishmanial drugs is far from satisfactory [9].

This demands the very requirement for effective drug discovery and development approach based on enzyme linked target identification; corresponding molecular pathway and cellular homeostasis of parasite involved in VL. Previous studies elucidated the role of various *Leishmania spp.* proteases involved in life cycle and pathogenesis of the parasite [10]. Among them zinc metalloprotease, Gp63 or leishmanolysin of *Leishmania spp.* have been identified as an important multifunctional virulence factor, found in abundance on the surface of *Leishmania spp.* promastigotes. It is also present in limited quantities inside the parasite [11] [12]. It is the key enzyme responsible for parasite propagation, promastigote binding to and internalization in macrophages as well as attenuation of formation of reactive oxygen intermediates favouring amastigote proliferation [13]. The myriad diversity as well as the high catalytic activity at body’s physiological temperature of this virulence factor favours the dissemination of the parasite in host [14] [15] [11] [16]. Apart from its major role in maintaining cellular homeostasis in parasite survival, recent experiments performed using discreet parasitic models demonstrated the protective nature of Gp63 against complement fixation and processing, which shields *Leishmania* promastigotes during its tarriance into mammalian hosts [17] [18] [19].

Considering the above scenario, we have designed a library of novel glycoside derivatives with D-glucose and N-acetyl-D-glucosamine as primary backbone template. We have used green chemistry approach which is a forward, unidirectional, single step synthesis process with no toxic byproducts and an unbiased method for designing target molecules [20]. Due to their volatile abundance in nature and relatively simpler conformation; glycoside derivatives are gaining current momentum as bioactive molecules in pharmaceutical industries [21].

These molecules were found to be completely non-toxic in mammalian cells and the hits showed promising result against both forms of *L. donovani* promastigotes as well as intra-macrophagic amastigotes. The Glycoside 2 **(Gly 2)** showed strong potency against lab and clinical strains of *L*. *donovani* at 1.13μM and 1.6μM (=IC50) concentration respectively, while mammalian cells were tolerant to the presence of **Gly 2** in mM range. **Gly 2** treatment leads to abrogation of parasite multiplication, induction of ROS generation and disruption of mitochondrial membrane potential leading to promastigote death. Besides its inhibitory role in *in-vitro* cultured parasites we have demonstrated its efficacy in body’s physiological condition where it marked prevention of parasite evasion from complement mediated lysis. Additionally, **Gly 2** was quite effective against the BS12, a clinical isolate of PKDL. Thus, owing to these observations, we evaluated the structure activity relationship (SAR) of the novel glycosides having hydrophilic head group and variable chain length of tail (lipophilic group). It helps to screen a large number of molecules spanning distinct regions of the bioactive chemical space; which is an inherent property seen in natural products [22]. We have demonstrated efficient binding of **Gly 2** molecule with Gp63 using Cellular Thermal Shift Assay (CETSA) and *in silico* studies. Furthermore, **Gly 2** displayed exemplary lethal effect on amastigote stage of parasite and found to be efficient enough in pathogen clearance within a macrophage infection model at 3.5μM (≥IC50). Additionally, **Gly 2** could target both VL and PKDL simultaneously. Thus, **Gly 2** not only proves to be absolute benign to mammalian cells but it also subverts the parasite mechanism of immune evasion making it as a potent antileishmanial against VL and PKDL.

## 2. Materials and methods

### 2.1 Parasite growth and maintenance

Promastigote-forms of *L. donovani* (Ag83 strain) were cultured at 26 °C in M199 media (GIBCO, India), pH 7.4 supplemented with 10% (v/v) inactivated Fetal Bovine Serum (FBS, GIBCO, India) and 0.02 mg/mL gentamycin (Life Technologies, USA). The clinical isolate of PKDL, BS12 was a gift from Prof. Mitali Chatterjee Lab (Institute of post graduate medical research, India). These isolates were routinely cultured at 22°C in M199 medium (GIBCO, India) with 100 U/mL penicillin-streptomycin (Gibco, Invitrogen, Thermo Fisher Scientific, NY), 8 μM hemin (4 mM stock made in 50% triethanolamine) (Sigma, USA), 25mM Hepes (N-[2-hydroxyethyl]piperazine-N0-[2-ethanesulfonic acid; Sigma]), supplemented with 10% heat inactivated FBS (GIBCO, India). Cultures were maintained between 10^6^ and 10^7^ cells/mL for continuous exponential growth. 1X10^6^ cells /mL parasite count was constantly maintained for all the experiments.

### 2.2 Cell Culture

The J774.A1 murine macrophage cells were grown in Roswell Park Memorial Institute (RPMI) 1640 Medium in the presence of 10% (v/v) FBS and penicillin-streptomycin (Gibco, Invitrogen, Thermo Fisher Scientific, NY) at 37°C (humidified) and 5% CO2. Primary MDCK cells were obtained from the National Centre for Cell Science, Pune, India. MDCK cells were derived from the kidney tissue of an adult female cocker spaniel. These were grown in Dulbecco’s modified Eagle’s minimal essential medium (DMEM) in the presence of 10% (v/v) FBS and penicillin-streptomycin (Gibco, Invitrogen, Thermo Fisher Scientific, NY) at 37°C (humidified) and 5% CO2.

### 2.3 General procedure for Synthesis of glycoside derivatives

Pre-stirred solution of glucose (200 mg, 1.11 mmol) or glucosamine (200 mg, 1.11 mmol) was prepared in neat alcohol (5-10 mL) and Amberlite IR 120-H^+^ resin (400 mg) was added to it. The resulting mixture was stirred at 100°C for 24 h. After completion of the reaction, the mixture was cooled down to room temperature, and filtered to remove the resin. The filtrate was evaporated under reduced pressure to obtain compound **1-3** as white solid in acceptable to good yield. Compounds **4**-**6** were prepared in good yield adopting similar reaction protocol using corresponding alcohol and glucose or glucosamine followed by acetylation using acetic anhydride in pyridine at room temperature. All the final compounds **1**-**6** were purified using flash column chromatography before using them for biological activity. The final compounds were formed as semi-solid or solid and were characterized by ^1^H-NMR and ^13^C-NMR. ^1^H-NMR and ^13^C-NMR spectra in CD3OD and D2O were recorded on Bruker 400 MHz spectrometer at ambient temperature. ^1^H recorded by 400 MHz and ^13^C recorded by 100 MHz. Proton chemical shifts are given in ppm relative to the internal standard (tetramethylsilane) or referenced relative to the solvent residual peaks (CD3OD: δ 3.31). Multiplicity was denoted as follows: s (singlet); d (doublet); t (triplet); q (quartet), m (multiplet): dd (doublet of doublet); dt (doublet of triplet); td (triplet of doublet); ddd (doublet of doublet of doublet), etc. Coupling constants (J) were reported in Hz. Column chromatography was performed by using silicagel 100-200 and 230-400 mesh. HRMS spectra determined from quadrapole/Q-TOF mass spectrometer with an ESI source (Supplementary text 1).

### 2.4 Treatment procedure

All the compounds were dissolved in dimethyl sulphoxide (DMSO) (Sigma-Aldrich) for 1mM stock solution and was screened against promastigotes at concentration of 5μM. For Glycosides (Gly) 2, 6 and 8 a range of concentrations starting from 100nM to 100μM were screened against promastigote form of parasite and mammalian cells; 5μM was the working concentration for these Compounds (>IC50 value) in all *in-vitro* experiments.

### 2.5 Anti-parasite drug susceptibility assay

Initially, promastigotes of Ag83 strain of *L. donovani* were exposed to various concentrations (100nM-100 μM) of each compound in a 96-well microtiter plate (100 μL/well volume) and were incubated for 72 h at 26°C. Parasites treated with amphotericin B (3μg/mL) (Sigma-Aldrich) were maintained as the positive control. Finally, LDH cytotoxic assay was performed as per standard protocol (CytoTox 96 Non-Radioactive Cytotoxicity Assay-Promega, USA). The percent cytotoxicity of test compounds was calculated by normalizing with amphotericin B as 100%. Colorimetric quantification of toxicity towards promastigotes upon treatment with various concentrations of the glycosides was carried out using LDH assay. Further percentage growth inhibition was calculated using the formula:

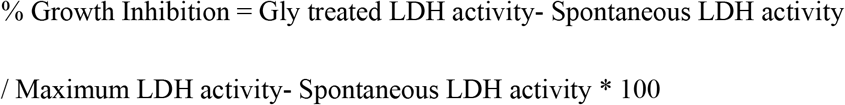

As per the formula, the spontaneous LDH Activity = activity of the untreated cells, and the maximum LDH activity = activity of the amphotericin B treated cells. These values were transferred in the GraphPad Prism version 8.0.1, and IC50 value for Gly 2, 6 and 8 treated was generated for Ag83 and PKDL strain of *L. donovani* using sigmoidal dose-response model with nonlinear regression tool.

### 2.6 Qualitative estimation of live /dead parasites

The toxic effect of the three compounds, Gly 2, 6 and 8 on promastigotes at 5μM > IC50 concentration was measured by Hoechst and PI double staining. After exposure with the compounds for 24 h, Parasites were harvested, PBS washed and stained with Hoechst 33258 (1μg/ml) (Life Technologies, USA) and PI (5 μg/mL) (Life Technologies, USA) followed by incubation for a period of 20min at 37°C. Subsequently, cells were washed for excessive stain removal and visualized using fluorescence microscope with 510-560 nm filter block for PI red fluorescence and excitation / emission spectra at 361/497 for Hoechst.

### 2.7 *In-silico* studies

The physicochemical properties of compounds were calculated using SwissADME [23] module provided in SIB (Swiss Institute of Bioinformatics) webserver (https://www.sib.swiss). The designed GLY compounds were evaluated for their ADME profile, including drug-likeness, partition coefficient, solubility, and oral bioavailability parameters according to Lipinski’s “rule-of-five”.

### 2.8 Antibody generation

Synthetic oligonucleotides encoding the catalytic motif of LdGp63 with PstI and HindIII overhangs were synthesized and ligated to PstI and HindIII digested pQELTB plasmid. [24], and transformed into *E. coli* DH5α cells for propagation and into *E. coli* M15 cells for recombinant fusion protein expression. *E.coli* M15 cells, containing the recombinant plasmid having the catalytic motif of LdGp63 in translational fusion with *E. coli* heat labile enterotoxin B subunit, were induced with 1 mM IPTG. Induction of expression with IPTG resulted in production of recombinant proteins as insoluble inclusion bodies. The recombinant fusion protein was purified to near homogeneity from solubilized inclusion bodies using metal affinity chromatography taking advantage of the histidine tag present at the N-terminal of the fusion protein. Female BALB/c mice were immunized with the purified fusion protein followed by a single booster on day 15 post immunization. Blood was obtained a week post booster administration through retro-orbital plexus. Anti-fusion protein antiserum was obtained after incubating the blood at RT for 1 h followed by centrifugation at 5000 rpm at 4 °C for 10 min. Antiserum thus prepared was stored at −20°C until further use.

### 2.9 Western blot analysis

Immunoblotting assay was performed using generated Mouse polyclonal anti-LdGp63 catalytic domain which were used at 1:500 dilution. The whole-parasite lysates were washed twice with 0.01M PBS, lysed in Lysis buffer (50mM Hepes, 150mM NaCl, 1% Triton X-100 and 1% IGEPAL, 1 mM phenylmethyl sulfonyl fluoride). These were denatured at 70°C. The samples were then homogenized with a 1-ml syringe and a 22-g needle before SDS-PAGE and blotting onto nitrocellulose membranes. The membranes were blocked with 5% skimmed milk powder dissolved in PBS containing 0.1% Tween-20.

### 2.11 Promastigote Proliferation Assay

Ag83 parasite growth and multiplication were assessed by Fluorescence Assorted Cell Sorting and fluorescence microscopy with 6-Carboxyfluorescein diacetate succinimidyl ester (CFDA-SE, Life Technologies, USA) as a probe. Promastigotes were washed thrice with 0.1M PBS. The cells were labelled with CFDA-SE dye and were then incubated at 37°C for 10 min during which they were mixed 3 to 4 times properly. These cells were then resuspended in ice-cold M199 medium. Further they were centrifuged at 1200g for 10 mins (4°C) and resuspended in fresh medium. Cells were treated with the Gly 2 and were analyzed through BD FACS DIVA for 3 consecutive replicates after 0, 24 h, 48 h and 72 h, respectively.

### 2.12 Estimation of ROS levels

The redox homeostasis of promastigotes was monitored both qualitatively and quantitively by 2-7 dichlorodihydro fluorescein diacetate (DCFDA) (Life Technologies, USA) staining. The untreated and Gly 2 treated promastigotes (1×10^6^ cells/mL) were cultured for 72 h. Following incubation, the parasites were harvested, washed with 0.1M PBS and stained with H2DCFDA (20μM) for 20min at 37°C. Excess stain was removed by washing with PBS and samples were resuspended in PBS (50μL) and followed by imaging. Fluorescence intensity was determined using an excitation filter at 485 nm and an emission filter at 535 nm using confocal microscope (Nikon Ti eclipse, USA). The samples were also analysed through BD FACS diva.

### 2.13 Quantification of mitochondrial membrane potential

Mitochondrial membrane (Δ*ψ*m) potential was examined using 5,6-dichloro-2-[3-(5,6-dichloro-1,3-diethyl-1,3-dihydro-2H-benzimidazol-2-ylidene)-1propenyl]-1,3-diethyl-,iodide (JC-1 dye) (Life Technologies, USA) as a probe. Treated and untreated groups were incubated for 24 h. Cells were washed with PBS, JC-1 labelled, and samples were analysed through FACS diva (Beckton-Dickinson, Pharmingen). The approximate fluorescence excitation/emission maxima of 514/529 nm for monomeric form and 585/590 nm for J-aggregate form were used. The labelled cells were also allowed to adhere to glass slides for visualization under Fluorescence microscope (Nikon Ti eclipse, USA); excitation and emission filters of TRITC and FITC were used.

### 2.14 Detection of Complement-Mediated Cytolysis of Promastigotes by uptake of Propidium Iodide

Parasites with an optimum 10% non-heat inactivated normal human serum were incubated with Gly 2 at 5μM concentration for 15 min. Promastigote lysis was detected by uptake of PI, by use of a FACS Diva flow cytometer (Beckton-Dickinson, Pharmingen), in accordance with the manufacturer’s protocol. Promastigotes were washed for excessive stain removal and filter block for PI red fluorescence, 510–560 nm was used.

### 2.15 *In-silico* membrane binding studies and docking studies

Using PerMM web server for theoretical assessment of passive permeability of molecules across the lipid bilayer, we checked the membrane binding studies with synthesized glucoside compounds. 3D protein structure of *L. major* GP63 (PDB ID-1lml) was obtained from Protein databank. BLASTp was used for LdGp63 to achieve sequence identity with template (*L. major* leishmanolysin in complex Zinc ion, 1LML). Homology modelling was further performed. RAMPAGE was utilized to obtain Ramachandran plot. Chemical structures of compounds were synthesized through the ChemSketch. Protein and Ligands structure were optimized using Swiss PDBviewer and ChemBioDraw ultra respectively. AutodockVina and Cygwin terminal were utilized to execute the docking commands. Binding site for the ligand was chosen around its catalytic domain residues. Chimera, Ligplus, Discovery Studio and Pymol softwares were used for further analysis of docking results.

### 2.16 Cellular Thermal Shift Assay

For a CETSA in promastigotes, parasites were seeded in 6-well cell culture plates (1.0 × 10^6^ cells /well) and exposed to Gly 2 at the 5μM >IC50 for 24 h in the incubator. Control cells were incubated with an equal volume of a vehicle. Following incubation, the parasites were washed with PBS to remove excess drug and harvested in 200μL of a lysis solution [50mM Hepes, 150mM NaCl, 1% Triton X-100 and 1% IGEPAL, 1 mM phenylmethyl sulfonyl fluoride]. The lysates were centrifuged at 14000g for 20 min at 4 °C, supernatants were transferred to new tubes. Further the protein concentration was measured using the Pierce BCA protein assay kit. After preparation of lysates, 30μL (0.8 mg/ml) aliquots of the supernatants were heated individually on a Thermomixer compact (Eppendorf) at different temperatures for 7 min and then cooled at room temperature for 3 min. CETSA samples were separated by sodium dodecyl sulfate–polyacrylamide gel electrophoresis, and immunoblotting was performed as described previously using a mouse polyclonal anti-Ld gp63 catalytic domain antibody (1:500).

### 2.18 Ethics statement

Normal Human Serum (NHS) is obtained from Rotary Blood Bank, Tughlakabad, New Delhi. All methods were carried out in accordance with relevant guidelines and regulations. Ethics Committees of Shiv Nadar University approved all the experiments carried out with NHS. The Animal Ethics Committee (IAEC Code #288/11) of the Jawaharlal Nehru University approved animal usage and procedures for generation of antibody. Ethical statement towards the clinical isolate of PKDL strain is available with host lab from where we have achieved the strain. The strain was cultured in BSL2 lab facility under guidelines institutional biosafety committee, Shiv Nadar University.

### 2.19 Statistical analysis

Student’s t-test was performed to evaluate significant differences between treatment and control samples in all the experiments performed using ANOVA test. P-value < 0.05 and <0.01 was considered to be significant indicated as * and ** respectively. Results represent the mean ± SD of minimum three independent experiments. Calculated IC50 value and all statistical analyses were performed by using GraphPad Prism version 8.01. Intensity profile for western blots were calculated using ImageJ software.

## 3. RESULT

### 3.1 Synthesis of glycoside-based compounds using a green synthetic route

The commercially available D-glucose 1 and N-Acetyl-D-glucosamine 2 molecules were coupled along with various short to long chain alcohols under acidic conditions, to design and synthesize glycoside-derived compounds. The synthesis of ethyl Glycoside *N*-Acetyl-D-glucosamine (Glycoside-1) was prepared by refluxing 1 in ethanol for 24 h in the presence of amberlite-H+ IR-120, which gave us the desired product Glycoside-1 as anomeric mixture in good yield as described **[Fig. 1A]**. The process does not involve any further purification and amberlite resin can be reused by activating it with 1 N HCl in MeOH. In parallel, other compounds in this series Glycoside-2 to Glycoside-6 were also prepared adopting similar reaction protocol (Methods), which resulted in good yields **(Table 1).** Furthermore, the alkyl Glycoside of N-Acetyl-D-Glucosamine Glycoside-7 to Glycoside-12 were prepared by refluxing N-Acetyl-D-Glucosamine with corresponding alkyl alcohols for 24 h in the presence of amberlite-H+ 100 IR-120 resin, which gave the desired products as anomeric mixture in good yield as shown **(Table 1).** Further characterizations of all 12 compounds were done using NMR and quadrupole/Q-TOF mass spectrometer with an ESI source determined using HRMS spectra. **(Supplementary text S1)**

**Figure 1:**
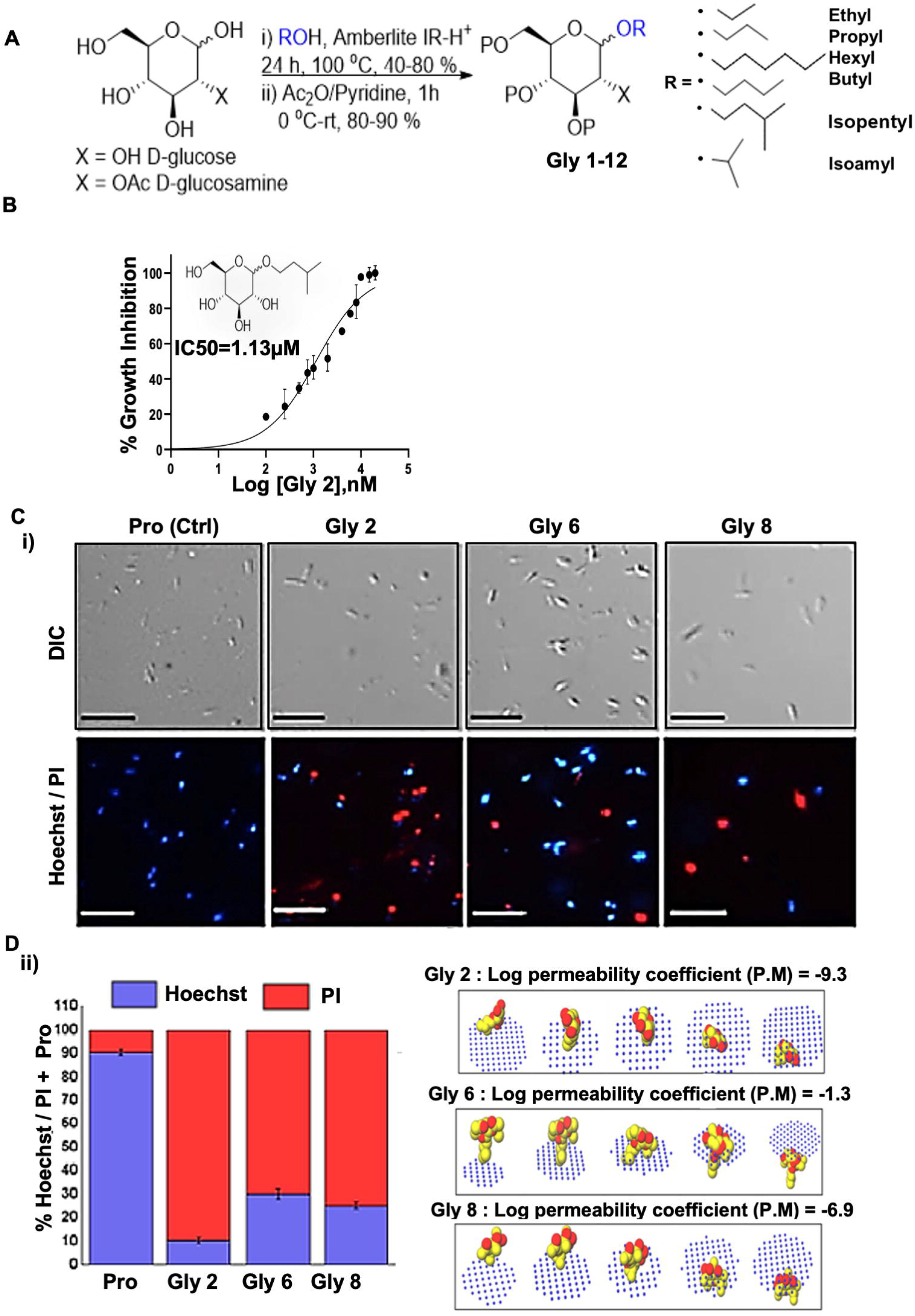
Scheme of synthesis of glycoside derivatives and concentration dependent effect of potent novel glycosides on promastigotes of Ag83 strain with variable membrane binding affinity. (A) Synthesis of designed glycosides was carried out with commercially available D-glucose and D-glucosamine coupling with various short to long chain alcohols under acidic condition; (B) Percentage inhibition of promastigotes treated with Gly 2 was evaluated using LDH Assay for 72 h and plotted as sigmoidal curve. Data normalization was done by taking into consideration the cytotoxicity induced by positive control (amphotericin B-3 μg/ml) as 100%. IC50 values for promastigotes of Ag83 strain were analysed using GraphPad Prism, represented as the mean ± SD where n=3, independent experiments; (C-i) the representative screenshots of PI stained promastigotes at 72 h assessed by fluorescence microscopy depicting parasite death on glycoside treatment; (C-ii) percentage Propidium Iodide (PI) positivity of the promastigotes at 72 h when treated by novel glycoside; (D) Membrane binding affinity of Glycosides 2,6 and 8 using PerMM server.

**Table 1:**
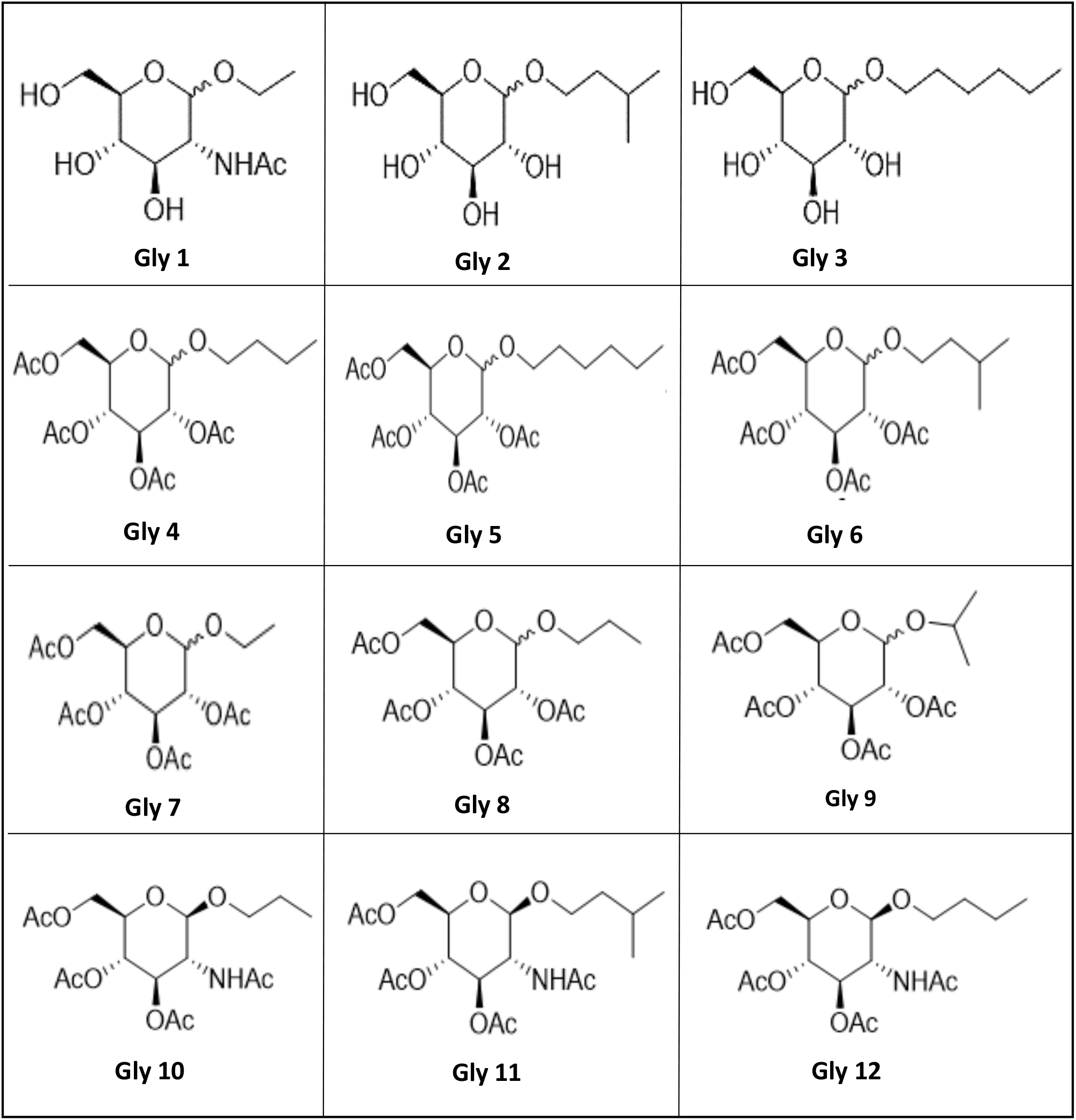
The structures of all the glycosides (1-12) are depicted.

### 3.2 ADMET and drug-likeness evaluation

SwissADME provides detail and extensive physicochemical profile, ADME, and medicinal chemistry property of a compound. The partition coefficient and solubility are the two parameters which play an important role in the physicochemical aspect. Based on predicted LogP value, it is concluded that all designed compounds lie within the range value of 1.6 to 3.6. All the 12 compounds depict the probability of a compound to be considered as drug candidate quality [25]. Even though LogP value does not always correspond to certain ADME aspect, this parameter could, where in this case .SwissADME LogP value was calculated from five different algorithm, therefore it is assumed that the value represents real condition [23].

### 3.3 *In-vitro* metabolic viability assay-based screening unravelled the lead glycoside derivatives and their toxic impact on *L. donovani* promastigotes

To evaluate the cytotoxic effects of 12 compounds on *L. donovani* promastigotes, Lactate Dehydrogenase Assay (LDH assay) was performed. This assay involves reduction of tetrazolium salts to formazan during LDH-mediated catalysis of lactate to pyruvate, which can be detected at 490 nM. Rate of formazan formation is proportional to the release of LDH through damaged cell membranes. At preliminary screening, the promastigotes were incubated with 12 compounds at a single concentration of 5μM respectively for a period of 72 h and the released LDH was estimated **(Supplementary Fig. 1A)**. Amphotericin B treated promastigotes were taken as positive control. Significant amount of formazan was observed in **Glycoside (Gly) 2, 6** and **8** treated samples suggesting that these glycosides could induce leakage of cytosolic LDH by exerting the toxicity on parasites. On incubation of promastigotes with **Gly 2, 6** and **8** compounds at various concentrations, the lethal effect of **Gly 2** (IC50=1.13μM) was found to be prominent than **Gly 6** (IC50=3.40μM) and **Gly 8** (IC50=1.61μM) comparatively **[Fig. 1B (i-iii)].**

As LDH release through damaged cell directly correspond to percentage of dead cells, this was further confirmed with live/dead dual staining of treated parasites with Hoechst and Propidium Iodide (PI) dyes. To achieve this, we treated promastigotes for a period of 72 h with **Gly 2, 6** and **8** and performed Hoechst/PI staining for fluorescent microscopy-based analysis. The results revealed that **[Fig. 1C]** untreated promastigotes showed viability (Hoechst+/PI-) whereas the treated parasites showed cellular death (Hoechst-/PI+). Ratio in the graph delineated the percentage viable/ dead parasites. Precisely, we found ~89.66% Gly2 treated parasites were dead as compared to ~70% and 75% in Gly 6 and Gly 8 treated promastigotes respectively **[Fig. 1C (i-ii)]**. We hardly found population of dead parasites in control **[Fig. 1C-i]**.

### 3.4 *In-silico* membrane permeability study revealed varied affinity of lead glycosides towards plasma membrane

Using Permeability of Molecules across Membranes (PerMM) server and database, we evaluated the passive permeability of small molecules across the lipid bilayer *in-silico.* We found that **Gly 2** was impermeable to the plasma membrane with log of permeability coefficient of −9.3 at temperature 310K and neutral pH of 7. On the other hand, **Gly 6** was permeable to plasma membrane with log of permeability coefficient of −1.3. In comparison of **Gly 2**, **Gly 8** was partially impermeable to membrane with log of permeability coefficient of −6.9. Further using GLMol the interactive 3D images of a compound moving across the membrane has been visualized **[Fig. 1D]**. This strengthened our hypothesis regarding probable interaction of **Gly 2** with membrane protein of parasite leading to suitable target identification. To evaluate this, we later performed *in-silico* docking studies with glycosides.

### 3.5 Lead Glycoside derivative (Gly 2) demonstrated minimal cytotoxicity in MDCK epithelial cells and J774A.1 macrophage

Minimum level of cytotoxicity could be detected for the compound in MDCK cells as well as macrophage cells. The CC50 values of **Gly 2** analogue was in the micromolar range. The selectivity index (SI) for the compound was calculated as the ratio between cytotoxicity (CC50) and activity (IC50) against promastigotes. The SI value for **Gly 2** were more than 1000 suggesting that the analogues are at least 1000 times more specific to parasites than host cells **(Supplementary Fig. 1C, D; Table 2).**

**Table 2:**
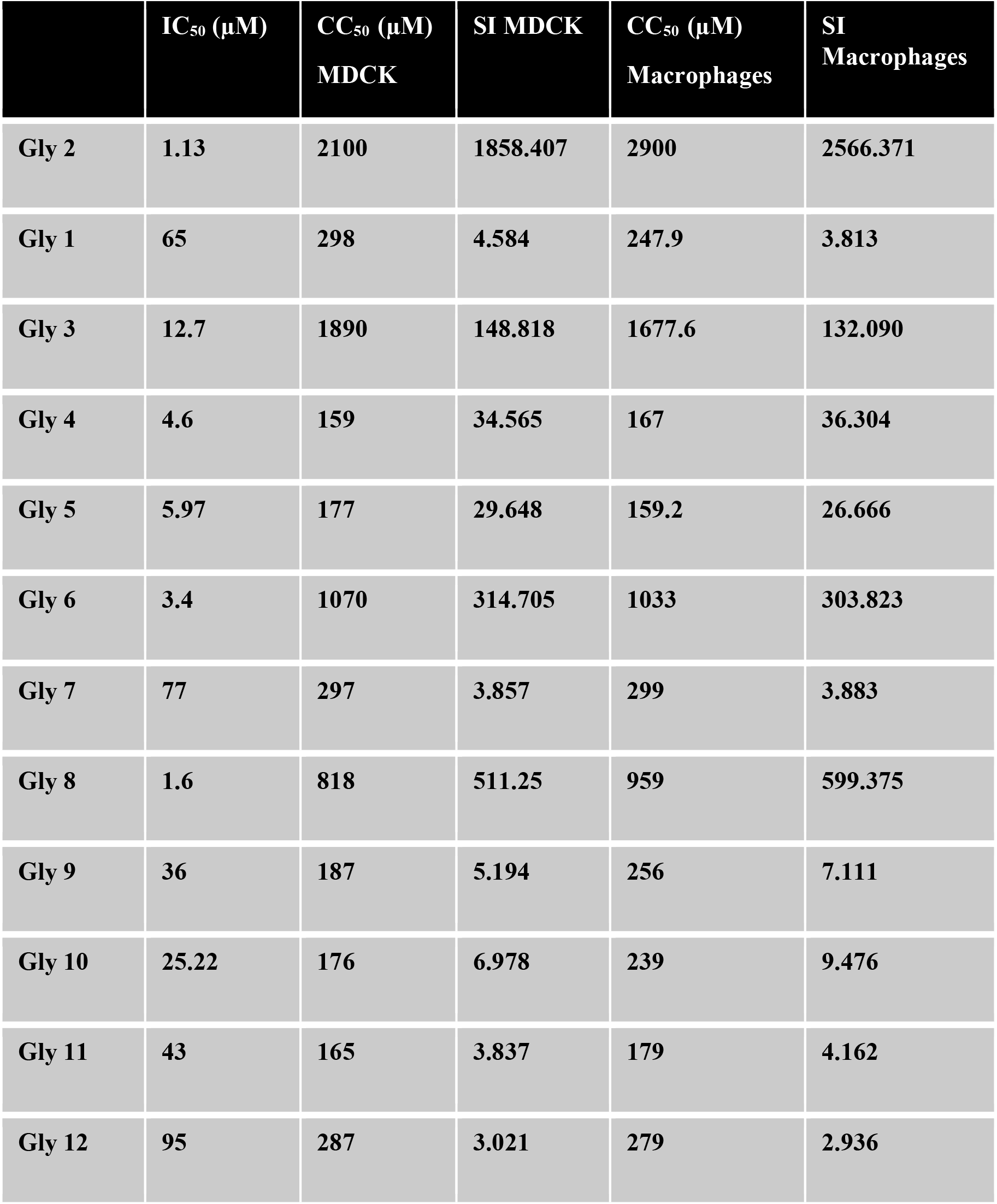
IC50/CC50/SI values of Gly 2

### 3.6 Generation of LdGp63 catalytic domain specific antibody

With the help of *In-silico* studies we found the amino acid residues involved in catalytic activity of Gp63 that pertaining to the catalytic domain (216-229 aa). Based on this, peptide containing catalytic domain was chemically synthesized which was tagged with LTB to increase the immunogenicity in mice. It was further injected in mice to develop polyclonal anti-LdGp63 antibody.

### 3.7 Treatment with Gly 2 abrogates promastigote proliferation

To further validate whether **Gly 2** treatment effectively hampers promastigotes multiplication or not; we assessed the proliferation of promastigotes by quantifying the release of cell permeable dye, CFDA-SE during cell division. This dye enters cells by simple diffusion, following cleavage by intracellular esterase enzymes to form reactive amine products, which covalently binds to intracellular lysine residues and other amine sources, producing detectable fluorescence. Decrease in fluorescence intensity is proportional to generation of promastigote daughter [26]. It was observed that the percentage of CFDA positive cells remained unchanged at both 24 h (81.59%), 48 h (74.68%) and 72 h (68.48%) at 5μM concentration of **Gly 2**, suggesting absence of cell division in the parental cell. However, the untreated promastigotes showed a reduced CFDA positivity 24 h, 48 h and 72 h suggesting cellular progression in a time dependent manner **[Fig. 2A-i].** Taken together, the results demonstrated a significant reduction in promastigotes’ growth and multiplication upon treatment with **Gly 2**. Further we also assessed the potential role of anti-gp63 antibody against promastigote proliferation. We found that there was subsequent change in percentage of CFDA positive cells in 24 h (70.70%) compared to 48 h (50.27%) and 72 h (39.29%) respectively **[Fig. 2A-ii]**. It clearly indicated that the raised polyclonal anti-Gp63 antibody could partially block promastigote proliferation for a short period of time. Thus **Gly 2** proves to be effective enough in controlling promastigote proliferation as compared to that of generated antibody.

**Figure 2:**
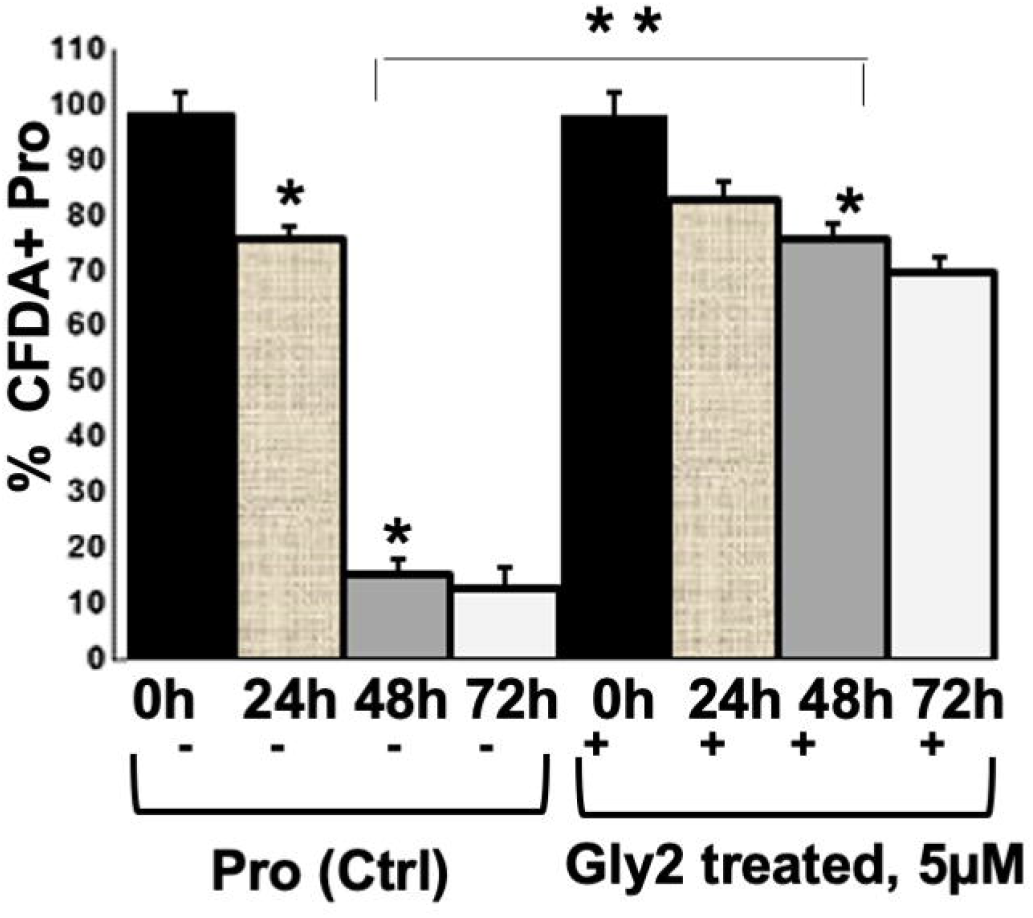
Effect of Gly 2 on promastigote proliferation using flow cytometry. Nominal decrement observed in percentage of CFDA-SE positive promastigotes when treated by compound for 24 h, 48 h and 72 h. The asterisks (**) indicate statistical significance (p < 0.01, n=3) between the indicated groups.

### 3.8 Anti-proliferative effect of Gly 2 led to destabilized redox potential in promastigotes

We investigated whether **Gly 2** mediated inhibition of LdGp63 disrupting multiplication could lead to destabilization of the redox potential in promastigotes by measuring ROS levels using 2’,7’–dichlorofluorescin diacetate (DCFDA) [27] [28]. Within cells, ROS oxidizes DCFDA into 2’,7’–dichlorofluorescin (DCF) which is a green fluorogenic compound. Intense green fluorescence in Gly 2 treated promastigotes indicate enhanced intracellular ROS generation, in treated parasites. The representative bar graph showed ~70% parasite population with DCFDA staining (DCFDA+/DCFDA-) indicative of elevated ROS levels **[Fig. 3A].**

**Figure 3:**
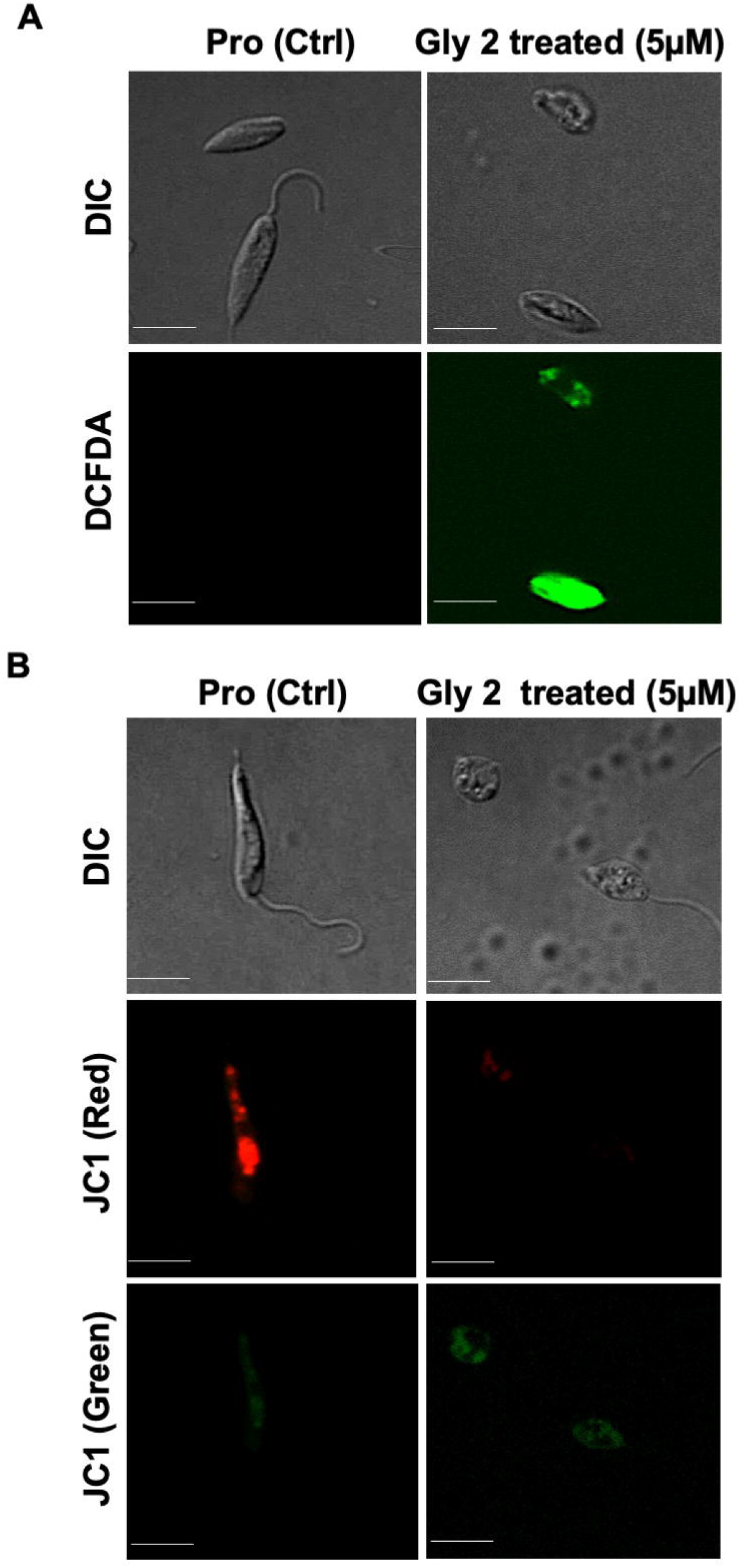
Gly 2 efficiently reduce promastigote proliferation activity leading to elevated ROS levels and disrupting mitochondrial membrane potential of glycoside treated promastigotes. A) Confocal imaging of DCFDA staining showed strong green fluorescence in Gly 2 treated cells, indicative of increased ROS levels. Presence of green fluorescence in treated parasites was observed in treated control, B) Effect of Gly 2 on ΔΨm of Ag83 promastigotes indicated by the conversion in monomer to oligomer forms of JC-1 using confocal micrographs. The shift in intensity of red fluorescence (JC1 aggregates/PE) to green fluorescence (JC1 monomers/FITC) implies ΔΨm in promastigotes following the treatment.

### 3.9 Gly2 interferes with mitochondrial function and disrupts mitochondrial membrane potential (ΔΨm)

Elevation of intracellular ROS coupled with the depolarization of mitochondria causing bioenergetics catastrophe is the hallmark of cellular necrosis [27] [29]. The maintenance of mitochondrial ΔΨm is vital for all metabolic processes and cell survival. To explore the effect of **Gly 2** on ΔΨm of promastigotes, we have used a lipophilic, cationic dye (JC-1) exhibiting green fluorescence, which enters the mitochondria and gets accumulated into a reversible complex known as J aggregates emitting red fluorescence [30].The fluorescence intensity was evaluated using confocal microscopy. Enhanced level of red fluorescence denotes more J aggregate formation due to higher ΔΨm, whereas shifting towards lower red or accumulation of higher green fluorescence implies a strong indication of destabilized ΔΨm. Notably, mitochondrial uptake of JC-1 dye was found to decrease with **Gly 2** treatment as compared to control healthy promastigotes which is then manifested as stronger green fluorescence due to monomeric JC-1 formation. Intense red fluorescence was observed in control groups suggesting JC-1 aggregation due to stable ΔΨm **[Fig. 3B].**

### 3.10 Gly 2 treated promastigotes enhances the complement mediated lysis in serum physiological condition

Mimicking the physiological condition of body, axenic promastigotes cultures in late log phase were exposed to complement-mediated lysis, which was determined by measuring propidium iodide (PI) stained cells [31] [32]. Histogram plots illustrate 50.02% population of promastigote death in 10% Normal human serum (NHS) within 30 min as compared to control which showed 0.06 % parasite death and 99.96% intact promastigotes in 10% FBS (heat inactivated). The **Gly 2** treated promastigotes however showed 79.05% parasite death in the presence of 10% NHS. There was 29.48% increment in PI uptake which is directly proportional to promastigote killing. Majority of **Gly 2** treated promastigotes were non-motile when observed under the microscope. The precise size of this population is difficult to calculate as, in the presence of NHS, promastigote cell volume and refractile properties are altered, blurring the distinction between promastigotes, cell debris, and NHS background signal. These results indicate that in the presence of **Gly 2** and 10% NHS causes rapid lysis Leishmania promastigote **[Fig. 4A].**

**Figure 4:**
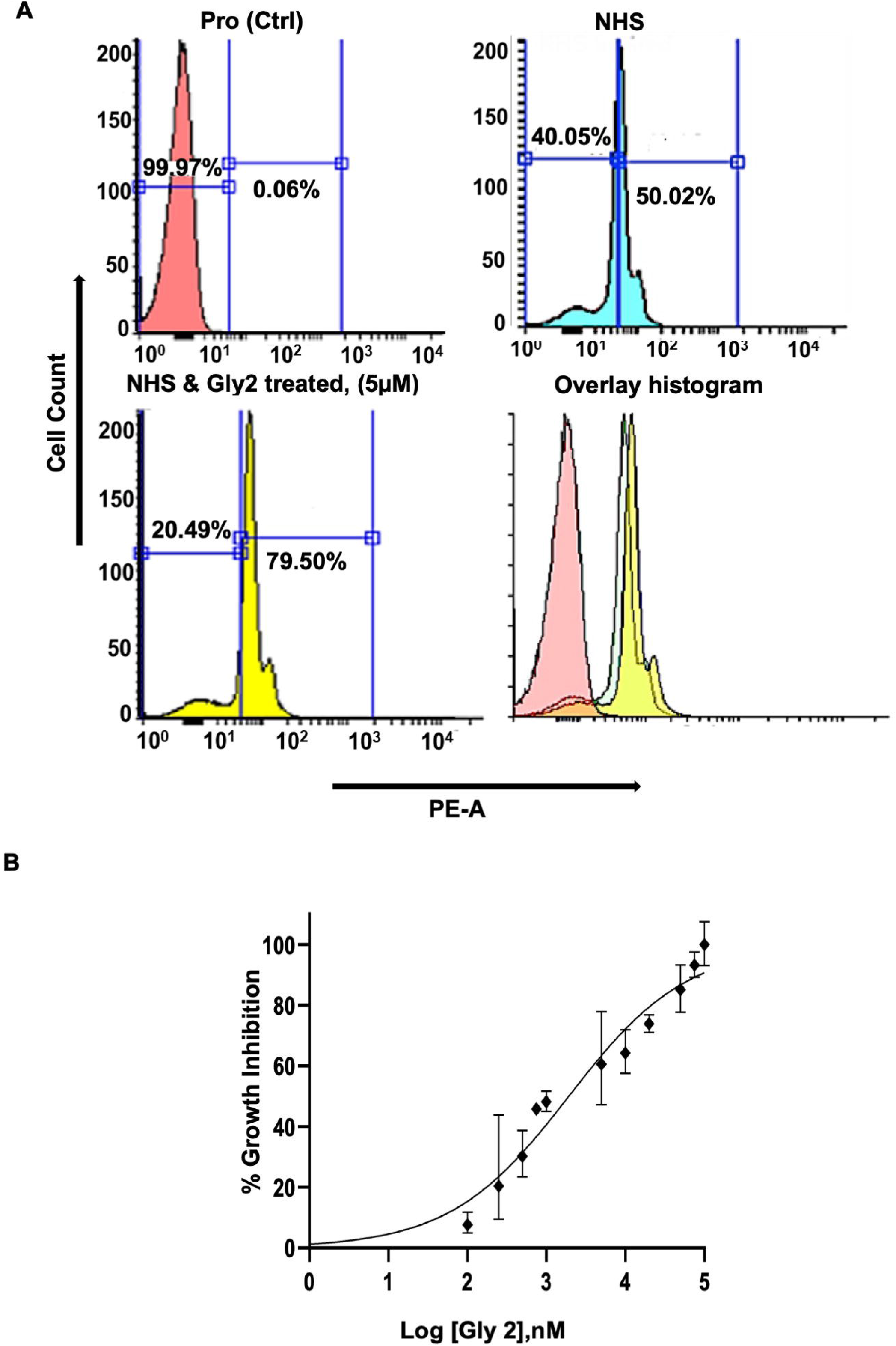
**Gly 2** treated promastigotes enhances the complement mediated lysis in serum physiological condition and its concentration dependent cytotoxic effect on PKDL strain. A) Promastigote lysis was analysed by uptake of PI using flowcytometry triggered by 10% Normal human serum; B) Cytotoxic effect of **Gly 2** on clinical PKDL isolate BS12 in concentration dependent manner.

### 3.11 Gly 2 inflicts cytotoxicity in Clinical PKDL strain BS12

We further evaluated the effect the **Gly 2** on the clinical isolate of PKDL strain, BS12 of *L. donovani* [33]. The results showed pronounced toxic effect of the **Gly 2** on the Indian origin leishmania isolate of PKDL. As mentioned previously, LDH assay was carried out on promastigotes treated with different concentrations of **Gly 2** for 72 h, while amphotericin B was used a positive control. The promastigotes showed IC50 value at 1.93 μM with 50% parasitic death **[Fig. 4B].** Along with Gly 2, we also evaluated the cytotoxic effects of 12 compounds on *L. donovani* promastigotes clinical isolate of PKDL strain at 5μM **(Supplementary Fig. 1B).** The data strongly suggests that **Gly2** could adversely affect the metabolic cell viability of clinical strain of PKDL.

### 2.12 *In-silico* docking of Gly 2 with 3D structure of *L. donovani-Gp63* (LdGp63) protein identified a druggable catalytic pocket bound to Gly 2

LdGp63 protein, a parasite receptor protein in its active form comprises of one chain containing multiple beta sheets and helices, consisting of 590 residues. Using BLASTp, we found that LdGp63 showed 80.9% sequence identity with template *(L. major* leishmanolysin in complex Zinc ion, 1LML). We performed homology modelling consequently with energy minimization for refined model of LdGp63 **[Fig. 5A & Supplementary Fig. 1E]**. Superimposition with this template rendered a RMSD value < 1 Å **[Fig. 5B].** Through RAMPAGE, we found that majority of residues were present in the favoured or allowed regions of the Ramachandran plot, thus validating the quality of the homology model **[Supplementary Fig. 1F]**. To evaluate the physical basis of protein-ligand interactions, we performed *in-silico* docking studies of LdGp63 with all compounds **(Supplementary Fig.2A)**. The best conformations of the docked compounds were selected based on their lowest free binding energy to the catalytic domain. Analysis of this pocket unravelled three residues (**His251, Glu252** and **Pro334**) that are involved in binding to the compounds **[Fig. 5C]. Gly 2** showed lower free binding energy as compared to others suggesting efficient docking **[Table 3].** In the docked complex, the O3, O4 and O5 residue of Gly 2 formed a strong H-bond interaction with **His251, Glu252** and **Pro334** residues of LdGP63 (2.83Å, 2.71 Å and 2.90 Å) respectively.

**Figure 5:**
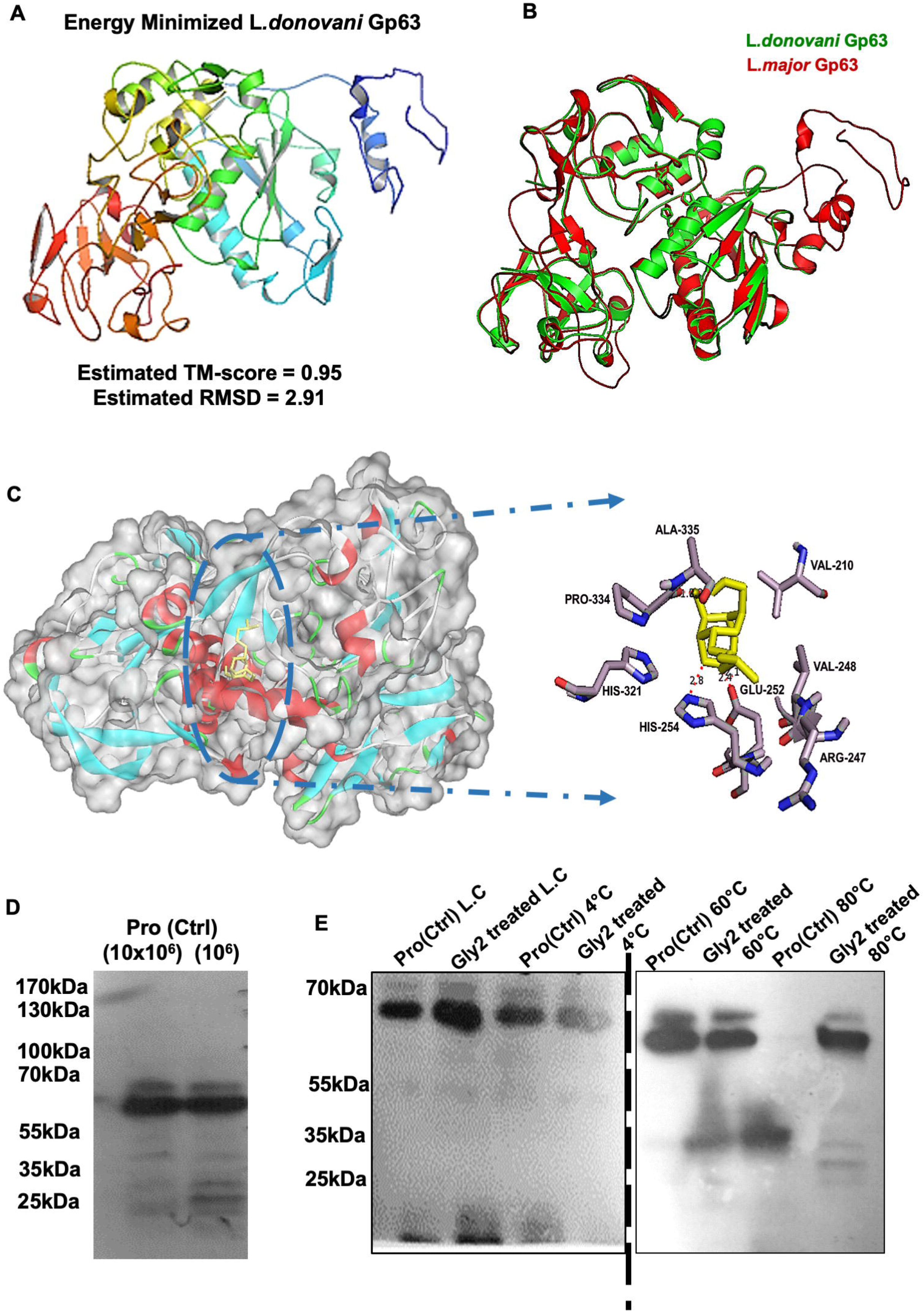
*In-silico* docking of Glycoside 2 **(Gly 2)** with 3D structure of *L. donovani-Gp63* (LdGp63) protein and *In-vitro* binding assay for potent LdGp63 inhibitor **(Gly 2)** in promastigote cell lysates using CETSA. A) RMSD value of LdGp63 model showed significant changes in stability post energy minimization; B) Superimposition of LdGp63 and LmGp63 models showed significant structure identity; C) 3D surface model of Gly 2-LdGp63 complex denoted by degree of hydrophobicity and surface accessible for ligand binding using a grey scheme. The defined region shows interacting residues of Gly 2-LdGp63 complex; D) LdGp63 was detected in variable number of promastigote lysate with generated polyclonal anti-LdGp63 antibody; cropped blots has been displayed, the full-length blots are included in a Supplementary Information figure; E) the thermostability of LdGp63 was analysed by immunoblotting following heat treatment at a range from 4 °C to 80 °C temperatures in treated or untreated promastigotes with Gly 2 (5μM) depicted as cropped blots from different gels.

**Figure-6:** Working model for discovery and translational application of **Gly 2.**

**Table 3:**
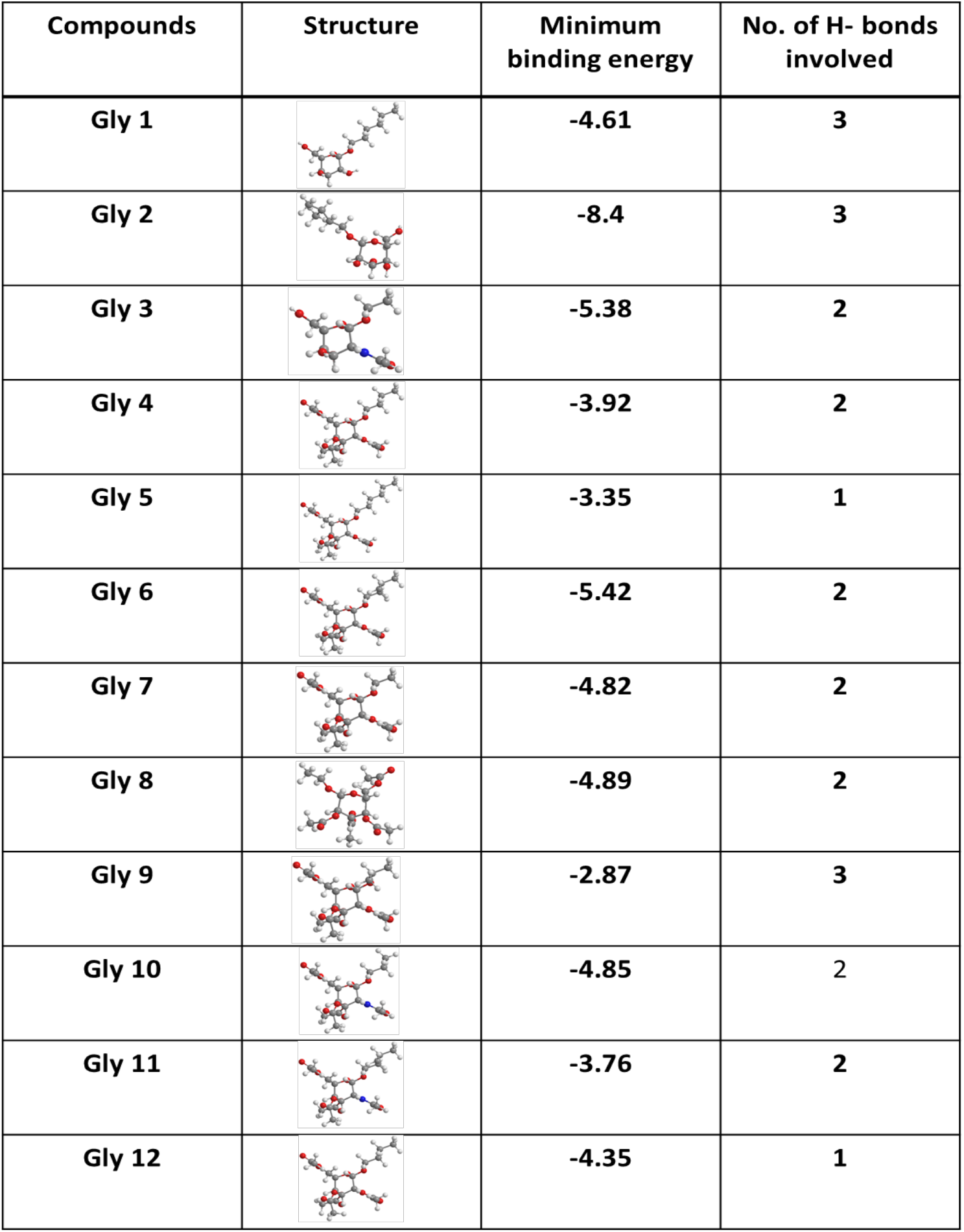
Critical residues lying within the docked ligand-protein complex were found to be forming H-bonds with highest minimum binding energy with Gly 2.

| Compounds | Structure | Minimum binding energy | No. of H- bonds involved |
| --- | --- | --- | --- |
| Gly 1 |  | -4.61 | 3 |
| Gly 2 |  | -8.4 | 3 |
| Gly 3 |  | -5.38 | 2 |
| Gly 4 |  | -3.92 | 2 |
| Gly 5 |  | -3.35 | 1 |
| Gly 6 |  | -5.42 | 2 |
| Gly 7 |  | -4.82 | 2 |
| Gly 8 |  | -4.89 | 2 |
| Gly 9 |  | -2.87 | 3 |
| Gly 10 |  | -4.85 | 2 |
| Gly 11 |  | -3.76 | 2 |
| Gly 12 |  | -4.35 | 1 |

### 3.13 Interaction of Gly 2 with LdGp63 enhances thermostability of protein-compound complex, detected using cellular thermal shift assay (CETSA)

Further to evaluate **Gly 2** dependent molecular mechanism involved in pathophysiological condition of parasite leading to its death; we verified the binding efficacy of Gp63, potent virulence factor with **Gly 2** inhibitor in promastigote lysates using cellular thermal shift assay **(CETSA)** [34]. The recently developed CETSA showed that target engagement can likewise be assessed in whole cell lysates based on altered protein thermostability [35], thus allows the investigation of target assignation under pathological conditions. We first titrated the number of promastigotes (10^4^-10^7^ cells) and immunoblotted with anti-Gp63 antibody **[Fig. 5D].** We then focused on treated and untreated promastigote lysates (10^7^ cells), which were incubated with **Gly 2** (5μM) or vehicle for 24 h and subsequently exposed to different temperatures before immunoblotting. The results of the immunoblot showed that the abundance of Gp63 Catalytic domain in the cell lysate decreased with increasing temperatures (4-80 °C). Importantly, in the range from 60 to 80 °C, the lysates treated with **Gly 2** showed abundances of Catalytic Domain of Gp63 protein higher than the abundances of the same protein in untreated. There was a drastic reduction in area intensity of bands in the promastigotes lysate at the temperature of 80°C which was measured to be 1842.062 as compared to Gly 2 treated promastigotes lysate which was indicated to be 20749.413 at similar conditions. The control lanes of 4°C constituting both untreated and Gly 2 treated parasite lysates exhibited area intensity of 20812.383 and 20755.839, respectively. This clearly suggested ligand-dependent stabilization by Gly 2 even at much higher temperature than the normal physiological temperature **[Fig. 5E].**

## 4. DISCUSSION

The major blockades towards effective treatment against leishmaniasis include pharmacokinetics and pharmacodynamics of drugs, toxicity, expensive, drug resistance, lack of vaccines and inadequate therapeutic interventions Successful treatment of cutaneous leishmaniasis with amphotericin B; a case of unresponsive to pentavalent antimony therapy [7]. The major prevention strategy involves protection of host from the bite of sand fly, which is an infeasible task. This creates a new pavement for discovering novel drug targets and developing target specific molecules; for better preventive measures and treatment modalities [36]. Recent unexplored area in the field of therapeutics against VL mainly focusses on transmission blocking drugs. Thus, targeting specific enzymes which are involved in cellular metabolism of promastigotes with transmission blocking drugs is an effective alternative strategy for efficient therapeutic implications [37].

Gp63/ Leishmanolysin belongs to a family of metalloproteinase which share the common signature sequence of HE*XX*H as a conserved zinc-binding/catalytic motif [38] [39]. It is present extracellularly on promastigote surface found in gut of the insect vector and intralysosomally parasitize macrophages as amastigotes in the mammalian host, causing human leishmaniasis. Gp63 has been suggested to function in multiple steps of the infection [40], i.e. their entry into macrophages [41] and intralysosomal survival [19] [42]. Gp63/Leishmanolysin thus appears to differ functionally from other HEXXH metalloproteinase families. Earlier reports elucidated the potent role of flavonoid glycosides from biologically active aqueous plant extracts against VL [43].

Based on these facts we have synthesized simple glucose and glucosamine backbone conferring natural product inspired library of Gp63 inhibitors using green chemistry approach. Upon screening of glycoside derivatives on *L. donovani* promastigotes, we found that **Gly 2, 6** and **8** imposed profound cytotoxic effects with no lethal effect in mammalian cells such as macrophages and canine epithelial cells, MDCK ensuring their specificity towards *L. donovani* promastigotes but not on host. The specificity was further assessed by their selectivity index ratio (CC50 for host cells/IC50 for parasite). The lead glycosides were more selective for *L. donovani* parasites than the host cells, with a selectivity index ratio of >1000. The low toxicity against mammalian cells is an important criterion while exploring new active compounds with anti-parasitic activity. The chemical structure of compounds is yet another essential factor related to their role as antileishmanials. The most active compound **Gly 2** derived from D-glucose have isopentyl as branched alkyl group with native free OH group which make the molecule hydrophilic (clogP 0.26) and less penetrable into cell. Whereas other active compound **Gly 6** having isopentyl as branched alkyl chain and hydroxyl group were masked with acetyl (OAc) group thus making molecule more hydrobhobic (clogP 2.4) suitable for cell permeability. The designed inhibitors were in fact supported by our *in-silico* experimental results where **Gly 2** is binding on the surface of the parasite *(Leishmania* promastigotes) whereas **Gly 6** easily penetrable through the cell surface. **Gly 8** were having shorter straight or linear alkyl chain (propyl) along with masked hydroxyl group making its amphiphilic (clogP 1.47) molecule that can perform dual role which includes surface binding probability and penetration inside the cell. We further validated the toxic effect of lead glycoside derivatives on promastigotes with Hoechst and PI staining leading to parasite death. We found that **Gly 2, 6** and **8** showed more PI+ and less Hoechst+ cells as compared to control promastigotes which depicted visa-versa. **Gly 2** showed highest PI+ among other two lead glycosides without induction of phospholipids exposure on the plasma membrane (data not provided).

To understand the cellular consequences of *in-vitro* inhibition of **Gly 2** promastigotes, we have investigated the promastigote proliferation. Using live staining with CFDA-SE (a strong membrane permeant dye), the effect of **Gly 2** treatment on promastigotes the growth kinetics of promastigotes was studied. The CFDA-SE dye upon cleavage by esterases within the cell, generates reactive amine products that covalently bonds with intracellular lysine to generate fluorescence. Based on this assay, the **Gly 2** treated promastigotes showed complete abrogation of cell division / progression at consecutive days. We even masked the promastigote surface with generated mouse polyclonal anti-LdGp63 catalytic domain antibody at a dilution of 1:400; interestingly found that such treatment lead to partial abrogation of promastigote proliferation.

Based on our proposition, **Gly 2** treatment lead to repudiation of promastigotes further causing increased ROS generation whereas control parasites showed balanced redox homeostasis. Further, it has been reported that drug-induced loss in ΔΨm is associated with cellular death in *L. donovani, L. amazonensis and Trypanosoma cruzi* [44]. ΔΨm loss and ROS elevation are coupled events and act as crucial signals of necrosis [45]. Thus, we then determined ΔΨm loss following increased load of oxidative stress in **Gly 2** treated promastigotes leading to necrosis like death which was corroborated with PI positivity proved earlier.

Later mimicking pathophysiological conditions of the body, we tried to elucidate the susceptibility of promastigotes towards complement mediated lysis. We propagated promastigotes in the presence of 10% NHS and after 15 min of incubation, 50% cellular death in control promastigotes was observed through incorporation of PI. In contrast to control parasites, there was 29% increment of parasite death in **Gly 2** treated samples, depicting 79% cytolysis.

Encouraged by these results, we evaluated the efficacy of **Gly 2** on clinical PKDL isolate BS12, which is responsible for the post kalazar disease manifestation, and known to play crucial role for *L. donovani* during interepidemic periods. It was observed that **Gly 2** leads to significant growth inhibition in BS12 at much lower concentration of 1.97μM>= IC50.

After confirmation of cytotoxic and downstream cellular effects on promastigotes of lab and clinical strains, we adopted an *in-silico* and *in-vitro* based Gp63 inhibition study. This was in accordance with previous reports suggesting role of Gp63 in promastigote multiplication [46]. The fundamental aim of the study was to ensure enhanced efficacy of the lead compound based on structure-to-activity relationships. Sequence and structure alignment between LmGp63 and LdGp63 show conserved Histidine triad as catalytic residues (His251, His254 and His320) in LDGp63. The *in-silico* docking study suggested that **Gly 2** is efficiently docking to Gp63 with low free binding energy, whereas docking with other compounds was inefficient. The **Gly 2** binding pocket represents an optimal volume (82 X 74 X 75 Å^3^) around catalytic domain for efficient binding to catalytic residues in the protein. Out of 12 molecules three of them (Gly 1, Gly 2, Gly 3) showed lower minimum binding energies such as −4.61, −8.4, −5.38 Kcal/mol respectively. However minimum binding energy of **Gly 2** is lowest as well as it interacts with the His251 residue present in the active site pocket of GP63. Among 12 glycosides, **Gly 2** shows strong hydrogen bond interaction with His251, Glu252 and Pro334. **Gly 2** also forms hydrophobic interactions around residues of LdGp63 catalytic pocket that assist it to fit in the catalytic pocket.

Further to assess strong interactions between a drug and its protein target in a physiologically relevant cellular environment we performed CETSA. Previous studies showed that CETSA has been performed in complex protein samples and in live cells displaying an altered melting or aggregation behaviour upon exposure to increased temperature [47]. CETSA has also been employed to identify *Plasmodium falciparum* protein targets in parasite lysates [48]. In this study, we have used target-specific antibodies that is antibody against catalytic domain Gp63. We found that on binding of **Gly 2** with Gp63; caused protein stabilization event under physiologic conditions even at remarkably high temperature of 60-80°C. Normally, at higher temperature protein denaturation takes place. But when the ligand is binding strongly to binding pocket of the protein; it affects both conformation and stability of the protein of interest. This was prominently observed when we immunoblotted the membrane with anti-LdGp63 catalytic domain antibody and found abundances of Catalytic Domain of Gp63 protein in treated samples. This suggested that there might be possibility that **Gly 2** has higher affinity in binding towards catalytic domain of Gp63.

Previously, there were reports suggesting that apoptotic parasites contribute towards macrophage invasion by balancing the ratio of viable to non-viable parasites as 1:1. However, our results demonstrated that more than 90% of **Gly 2** treated parasites are undergoing cellular death. Therefore, we hypothesize that the treatment with the **Gly 2** molecule will significantly reduce the number of parasites available for infection in macrophages. The results demonstrated potential effect on intracellular amastigotes at nanomolar range suggesting their strong anti-leishmanial activity in macrophage infection model. Details of the strategy have been explained in the working model.

## 5. CONCLUSIONS

In current scenario there is an estimation of 0·7–1 million newly reported cases of leishmaniasis erupting out every year from the 100 endemic countries globally. Thus, there is a requirement to develop new potent antileishmanial which are less prone to resistance development. Conclusively, we have introduced a novel glycoside derivative, Gly 2 as potential anti-leishmanial and suggested a prospective first in class therapeutic measure for both Kala azar and post kala azar dermal (PKDL) leishmaniasis respectively to ascertain their clinical utility.

## Acknowledgements

AC and NJ are supported by Shiv Nadar Foundation fellowships. Center for Informatics located within Shiv Nadar University is also duly acknowledged. SG is thankful for the funding support from DST INSPIRE grant. SP is grateful for the funding support from Shiv Nadar foundation. CN acknowledges the fellowship supported by CSIR-NET. We sincerely acknowledge help from Prof. Madhubala Rentala, JNU for gifting the J774.A1 murine macrophage cell line.

